# The positive inside rule is stronger when followed by a transmembrane helix

**DOI:** 10.1101/014175

**Authors:** Minttu T. Virkki, Christoph Peters, Daniel Nilsson, Therese Sörensen, Susana Cristobal, Björn Wallner, Arne Elofsson

## Abstract

The translocon recognizes transmembrane helices with sufficient level of hydrophobicity and inserts them into the membrane. However, sometimes less hydrophobic helices are also recognized. Positive inside rule, orientational preferences of and specific interactions with neighboring helices have been shown to aid in the recognition of these helices, at least in artificial systems. To better understand how the translocon inserts marginally hydrophobic helices, we studied three naturally occurring marginally hydrophobic helices, which were previously shown to require the subsequent helix for efficient translocon recognition. We find no evidence for specific interactions when we scan all residues in the subsequent helices. Instead, we identify arginines located at the N-terminal part of the subsequent helices that are crucial for the recognition of the marginally hydrophobic transmembrane helices, indicating that the positive inside rule is important. However, in two of the constructs these arginines do not aid in the recognition without the rest of the subsequent helix, i.e. the positive inside rule alone is not sufficient. Instead, the improved recognition of marginally hydrophobic helices can here be explained as follows; the positive inside rule provides an orientational preference of the subsequent helix, which in turn allows the marginally hydrophobic helix to be inserted, i.e. the effect of the positive inside rule is stronger if positively charged residues are followed by a transmembrane helix. Such a mechanism can obviously not aid C-terminal helices and consequently we find that the terminal helices in multi-spanning membrane proteins are more hydrophobic than internal helices.

## 1 Abbreviations

- mTMH - Marginally hydrophobic transmembrane helix.
- *TMH*_*sub*_ - Subsequent helix to the mTMH
- *TMH*_*pre*_ - Preceding helix to the mTMH

## 2 Introduction

The vast majority of transmembrane *α*-helical proteins are integrated into the membrane co-translationally via the Sec-translocon machinery. While the major determinant for membrane integration is hydrophobicity^1,2^, many multi-spanning membrane proteins contain transmembrane segments of surprisingly low hydrophobicity^3^. The membrane integration of such marginally hydrophobic transmembrane helices (mTMHs) can depend on sequence features outside of the hydrophobic segment itself, including the positive inside rule, flanking residues and interactions with neighboring helices^3–8^.

Positive charges in cytoplasmic loops have been shown to compensate by ≈0.5 kcal/mol towards the hydrophobicity of a given transmembrane helix^9^. This was also noted in our earlier study, where inclusion of adjacent loops strongly increased the insertion of five out of sixteen mTMHs^3^. In four of these, more positively charged residues were included in the cytosolic flanking loops than in the lumenal loops, see Table 1. However, flanks did not improve insertion for the other nine mTMHs, although in most cases the cytosolic loops contained more positively charged residues than the outside loops, see Table 1.

**Table 1:**
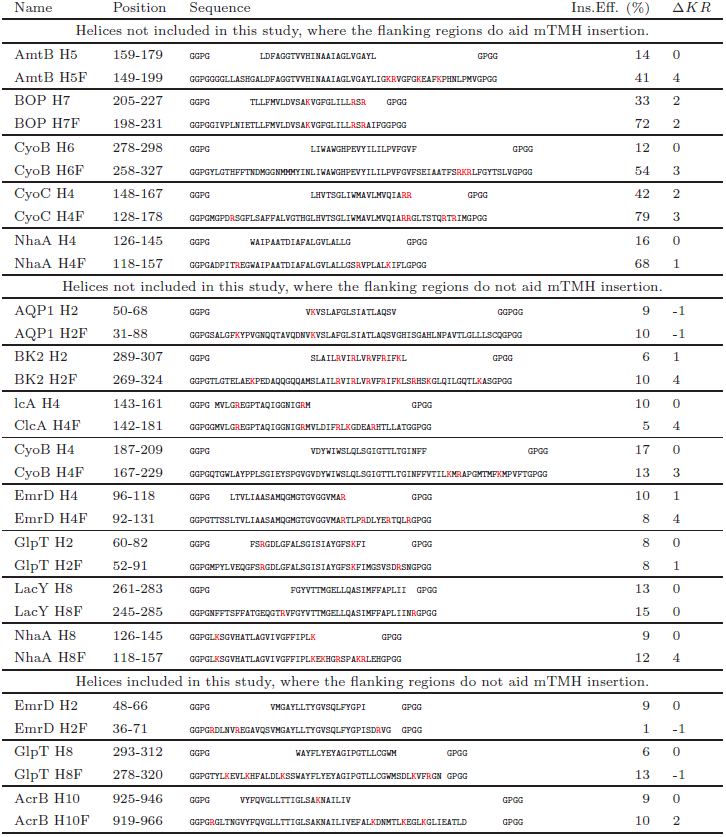
Translocon recognition efficiency (Ins.Eff.) of marginally hydrophobic helices with and without “flanking” regions (marked with F) from our earlier study^3^. The sequences tested are shown, followed by the experimentally measured insertion efficiency and the difference between positively charged residues in inside and outside loops (Δ*KR*). All positively charged residues are marked in red. The upper panel of the table shows the sequence where the flanks aid in the recognition. In all but one case the flanks adds positive charges on the loop following the mTMH. The second panels shows constructs where flanks alone are not sufficient to ensure integration into the membrane. Also here, in most cases the positive inside effect would favor the insertion when the flanks are included. The third panel shows the constructs tested here. For EmrD and GlpT the inclusion of flanks increases the number of positive charges on the outside loop, which causes the mTMH to insert even worse. In AcrB the positive inside effect is strengthen by the flanking residues but insertion is not improved.

Specific interactions between polar residues have been shown to reduce the overall cost of integrating a marginally hydrophobic helix to the membrane ^7^,^8^,^10^. In an earlier study, we found three naturally occurring marginally hydrophobic helices (EmrD mTMH2, GlpT mTMH8 and AcrB mTMH10), which all need the presence of their subsequent helix for efficient translocon recognition^3^.

To identify interactions between neighboring transmembrane helices during the translocon recognition of marginally hydrophobic helices, substitution scans with alanine/isoleucine were performed on the subsequent helix. Surprisingly, we did not find any evidence for specific interactions in any the three helices tested. Residues found to influence the insertion, were positively charged arginines at the N-terminus of subsequent helices in EmrD and GlpT. However, inclusion of these arginines alone, in the absence of the rest of the subsequent helices, does not aid insertion. Hence, the positive inside effect of these residues by themselves is not sufficient to aid the insertion of the marginally hydrophobic helix. Instead, they contribute to an orientational preference of the subsequent helix, which in turn lowers the apparent hydrophobicity barrier for the marginally hydrophobic helix and allows it to insert into the membrane^9^. Supporting the importance of this effect, we note that the terminal transmembrane helices are rarely marginally hydrophobic.

## 3 Results

All experiments in this study are performed using a previously described glycosylation assay^1–3^. The marginally hydrophobic helices (mTMHs) together with adjacent loops and neighboring helices are cloned into leader peptidase (Lep) as “H-segments”. When expressed *in vitro* in the presence of dog pancreatic microsomes, the topology of the constructs can be determined based on the number of attached glycans, see Figure 1 and materials and methods. For studies of the mTMHs and their subsequent helices (*TMH*_*sub*_) in EmrD and GlpT the Lep-I system is used, i.e. these helices are placed after the transmembrane helices of Lep. In AcrB it is also necessary to include the preceding helix (*TMH*_*pre*_) for efficient translocon recognition and therefore the Lep-II system is used. Here, the construct replaces the second transmembrane helix in Lep.

**Figure 1:**
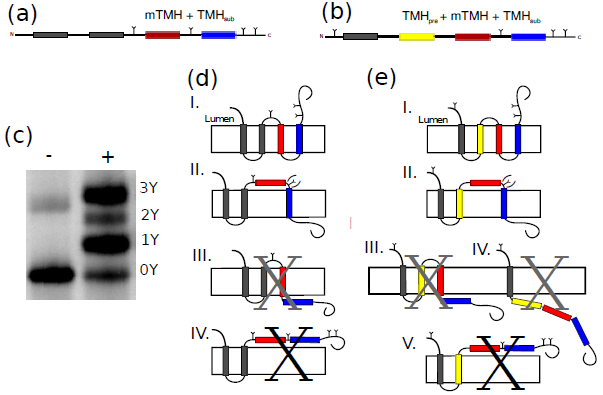
*In vitro* expression of leader peptidase (Lep) hosting mTMH segments consisting of mTMHs, adjacent loops and neighboring TMHs. The constructs are placed within Lep (in gray). a) For EmrD and GlpT the mTMH (in red) and the *TMH*_*sub*_ (in blue) are placed within the P2 domain of Lep. b) AcrB *TMH*_*pre*_ (in yellow), mTMH and the *TMH*_*sub*_ replace the second Lep TMH. In all constructs four engineered N-linked glycosylation sites (Y) are introduced to allow topology determination, since glycosylation only occurs in microsome lumen. c) By separating the *in vitro* expressed constructs on SDS-PAGE, the fraction of protein with different number of attached glycans can be quantified and their topologies determined^16^. Here EmrD mTMH2+TMH3 in LepI is shown expressed without (-) and with (+) microsomes. Bands appearing on SDS-PAGE are labeled according to how many glycans are attached. d) When the mTMH, along with *TMH*_*sub*_, from GlpT or EmrD is integrated into the membrane. e) For AcrB, the three-glycan form dominates when the mTMH and both *TMH*_*pre*_ and *TMH*_*sub*_ are inserted. Singly, doubly (or for AcrB quadruply) glycosylated forms all represent topologies where mTMH is not properly inserted into the membrane for all constructs. Topologies corresponding to differently glycosylated species are all drawn. Those we do not observe at all, or only very rarely, are crossed over in black. Those we deemed as unlikely due to hydrophobicities of the TMHs are crossed over in grey, see methods.

Inclusion of flanking residues did not improve the insertion of mTMHs in EmrD or GlpT, see Table 1. However, the presence of a hydrophobic *TMH*_*sub*_ improved both, see Figure 2. Efficient insertion of the mTMH in AcrB requires that also the TMH_pre_ is included, see Figure 2. We speculated that particular interactions between the mTMH and the subsequent or preceding helix might be of importance for the insertion^3^. As can be seen from the crystal structures, the mTMH in AcrB makes contacts with both the preceding and subsequent helices, while the mTMHs of EmrD and GlpT only interact weakly with the neighboring helices, see Figure 2. However, this lack of interactions not rule out the existence of important transient interactions between the helices during insertion and folding. By systematically replacing all residues in the subsequent helices, we tried to identify residues that interact with the mTMHs during the translocon recognition. Residues within the mTMH itself are not changed, i.e. its hydrophobicity remains identical in all constructs. In GlpT the scan was performed using alanine substitutions whereas isoleucine was used in EmrD and AcrB. The reason for not using alanine in EmrD and AcrB is that it caused many of the constructs to be cleaved by signal peptidase, resulting in ambiguous results.

**Figure 2:**
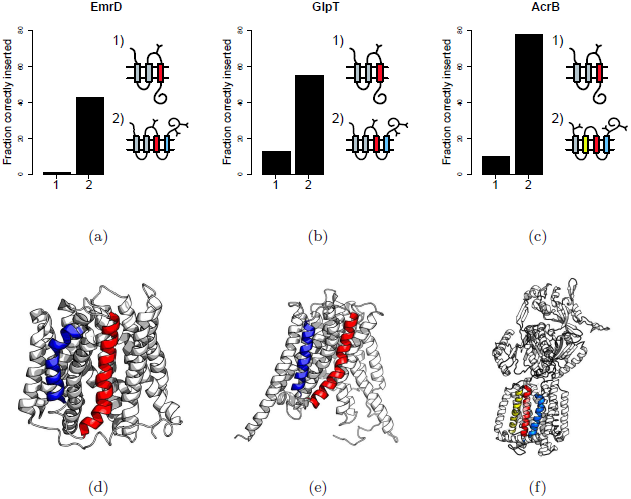
Overview of the three marginally hydrophobic helices in this study. a) The mTMH in EmD is not recognized by the translocon in the absence of
the subsequent helix. The presence of *TMH*_*sub*_ increases the insertion from 10% to 43%. b) GlpT mTMH is poorly recognized by the translocon and only 13% insert into the membrane. When *TMH*_*sub*_ is included, the membrane integration increases up to 55%. c) In AcrB the mTMH alone is inserted to 9% but inclusion of *TMH*_*pre*_ and *TMH*_*sub*_ improve the recognition to 86%). d-f) Structural representations of the three proteins; In all proteins the mTMHs are colored in red, *TMH*_*sub*_ in blue and *TMH*_*pre*_ in yellow. The helices are not in close proximity in EmrD and GlpT but do make contacts in AcrB.

### 3.1 N-terminal arginines in TMH_sub_ aid the insertion of the marginally hydrophobic helices

Some common features are shared by the local sequence context of the three mTMHs studied here; all loops between mTMH and *TMH*_*sub*_ as well as the N-termini of subsequent helices contain positively charged residues. These residues will reside on the cytoplasmic side given that the mTMH is properly inserted into the membrane, i.e. they contribute to the positive inside effect.

The inclusion of *TMH*_*sub*_ improves the marginally hydrophobic helix insertion in EmrD from about 10% to almost 50%, see Table 1. When either of the arginines, located at the N-terminal of the *TMH*_*sub*_, are substituted with isoleucine the mTMH is not inserted efficiently, see Figure 3. Except for P74I, where the substitution increased insertion efficiency, all other substitutions have only marginal effect on the insertion of the mTMH.

**Figure 3:**
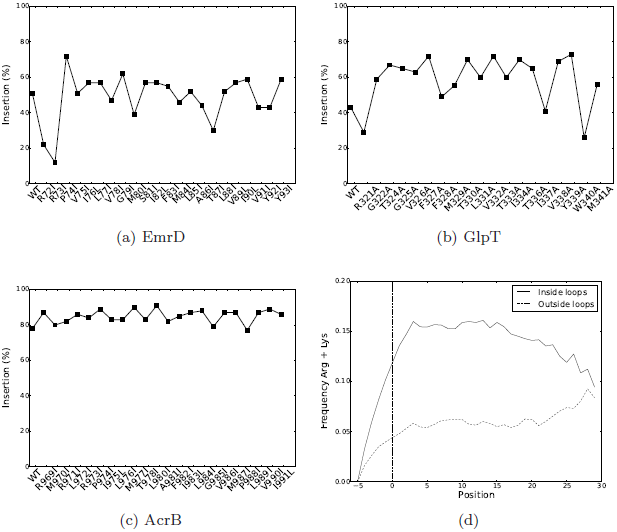
(a-c) Single substitution scans of *TMH*_*sub*_ in EmrD, GlpT and AcrB showing the average of two independent replicates. d) The frequency of positively charged residues in “inside” and “outside” loops given the distance from the predicted membrane termini.

In GlpT the mTMH with flanks is only inserted to 8%, but the addition of *TMH*_*sub*_ brings the membrane integration up to 43%. Much like in EmrD, substituting the N-terminal R321 with an alanine decreases insertion, see Figure 3B. One additional residue was detected in the alanine scan, the C-terminal tryptophan W340. Here, an alanine substitution reduces the insertion of the mTMH to 26%.

The marginally hydrophobic helix in AcrB requires both the subsequent and preceding helix to be efficiently recognized (86%). Similarly for the other two constructs, most isoleucine substitutions did not influence membrane insertion of the mTMH. However, in contrast to EmrD and GlpT, none of the arginines at the N-termini of *TMH*_*sub*_ are sensitive to isoleucine substitutions, see Figure 3C. Further, double and even triple isoleucine mutants are still able to enhance helix 10 membrane integration, see Figure 4.

**Figure 4:**
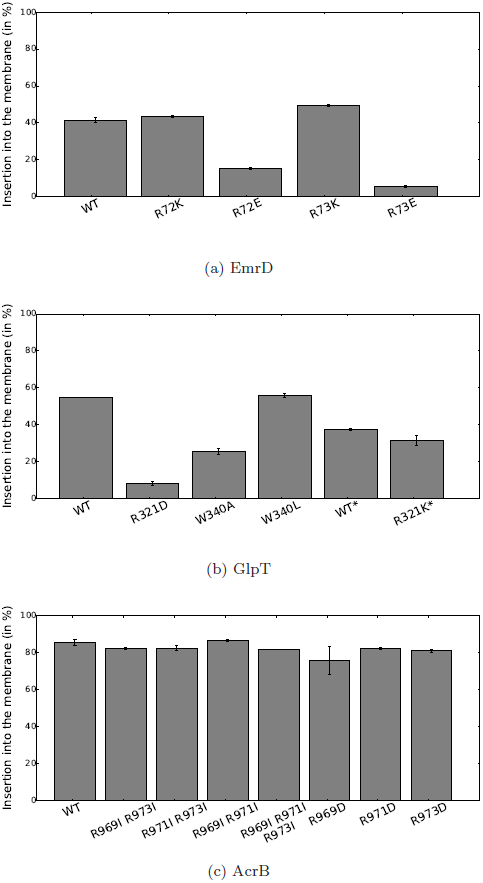
Replacement of positively charged residues at the N-terminus of *TMH*_*sub*_ in EmrD, GlpT and AcrB. a) EmrD R72 and R73 were substituted with the lysine or glutamic acid. Here, the experiments were performed using a new batch of microsomes, in which the native sequence insert up to 42% compared to 51% for the old batch. b) GlpT R321 and W340 replacements. Here, R321K substitution was done using a 22ew batch of microsomes and should be compared towards the insertion efficiency of the construct with native sequence expressed in the new microsomes (both marked with ^*^). c) Double and triple arginine to isoleucine substitutions for AcrB *TMH*_*sub*_. All experiments reported here were carried out using the new microsome batch. For AcrB the membrane integration was 86% in the new batch compared to 78% in the old batch. In all plots error bar reflects the spread of two independent replicates.

In EmrD and GlpT arginines can be successfully replaced by positively charged lysines but neither aspartic acids nor glutamic acids are tolerated, see Figure 4. Also, when W340 is replaced by a leucine the membrane integration of the marginally hydrophobic helix in GlpT is still enhanced, indicating that the hydrophobicity of the subsequent helix is important, rather than specific interactions made by W340. Conclusively, positively charged arginines are responsible for the membrane integration of marginally hydrophobic helices in EmrD and GlpT, while in AcrB they are not. One possible explanation for this is the presence of three lysines in the inside loop of AcrB causing a stronger positive inside effect, see Table 1.

Additional support for the importance of positively charged residues at the termini of helices is provided by bioinformatic analysis of alpha-helical membrane proteins, see Figure 3D. In the center of a transmembrane helix positively charged residues are virtually non-existent, but approximately five residues from the termini they begin to appear. The frequency then increases to reach a plateau a few residues outside the membrane. The frequency of positively charged residues in inside loops is approximately 15% compared to 5% in outside loops. About thirty residues away from the membrane the two frequencies approach the background frequency of arginine/lysine in non-membrane proteins (11.7%).

### 3.2 Orientational preferences of subsequent helices are responsible for the insertion of the marginally hydrophobic helices in EmrD and GlpT

Positive charges can lower the hydrophobicity threshold for an mTMH either directly or by contributing to an orientational preference of neighboring helices, i.e. in this study by the *TMH*_*sub*_. Positively charged residues on the cytoplasmic side of the membrane primarily cause both effects. However, to distinguish between orientational preference and the positive inside effect, it is necessary that the positively charged residues are located within the transmembrane helix and that they do not affect the orientational preference of the neighboring helix in the absence of the rest of the helix. The later requirement is necessary as the exact ends of transmembrane regions are difficult to predict^11^ and they might shift during the biogenesis of the protein^12^.

Here, we distinguish between the direct positive inside rule and orientational preferences of transmembrane helices by truncating the subsequent helix after the arginines located at the N-terminal parts of these helices, see Figure 5. In EmrD and GlpT the presences of only the arginines respective arginine from *TMH*_*sub*_ do not suffice for the recognition of the mTMHs. However, in AcrB the mTMH is efficiently integrated into the membrane even when *TMH*_*sub*_ is truncated after the third arginine. Hence, we conclude that the insertion of mTMHs in EmrD and GlpT is dependent on the orientational preference of their respective *TMH*_*sub*_, while for AcrB the positive inside rule is sufficient. However, in AcrB the inclusion of *TMH*_*sub*_ with the arginines replaced is also sufficient for efficient recognition. Hence, efficient insertion of the mTMH in AcrB is either dependent on the presence of *TMH*_*sub*_ or additional positive charges in the subsequent loop. The difference is most likely caused by the loop in AcrB, which provides a much stronger positive inside bias than in EmrD and GlpT, see Table 1.

**Figure 5:**
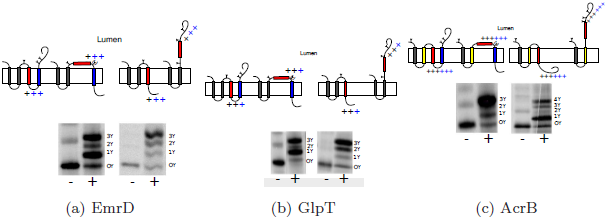
Cartoons of the full-length and truncated constructs are shown on the top of representative gel images of in vitro expressed constructs without (-) and with (+) microsomes. The modified glycosylation sites in each case are indicated with an Y and the number of glycans corresponding to each band on SDS-PAGE are indicated. The loop between mTMH and *TMH*_*sub*_ is short and can not be efficiently glycosylated, hence placed within parenthesis. The positive charges in the loop between mTMH and *TMH*_*sub*_ are shown in black whereas the arginine residues belonging to *TMH*_*sub*_ are shown in blue. a) EmrD mTMH2 together with *TMH*_*sub*_ is integrated to 40%, indicated by a thick band corresponding to the 3Y form. When truncated after R73 the mTMH is only inserted to 8%. b) When GlpT *TMH*_*sub*_ together with the first *TMH*_*sub*_ insert to 38%. When only residue R321 from *TMH*_*sub*_ is included, only 8% is inserted. c) For AcrB both the construct where mTMH and *TMH*_*sub*_ are present as well as one where *TMH*_*sub*_ is truncated after R973 insert well, 86% respetive 80%.

## 4 Discussion

Many membrane proteins contain marginally hydrophobic helices that are not independently recognized by the translocon^3^. How these helices are integrated into the membrane is not well understood. At least three mechanisms to aid insertion have been proposed, (i) the positive inside rule^13^, (ii) orientational effects caused by neighboring helices^9^ and (iii) specific interactions between two adjacent transmembrane helices^7^. In order to identify positions responsible for the increased translocon recognition we studied the effects of point mutations in subsequent helices of three natural marginally hydrophobic helices.

In summary, none of the marginally hydrophobic helices studied here were dependent on specific interactions with the subsequent helix. Positive charges on the other hand were found to be important by causing an orientational preference for the subsequent helix.

In EmrD and GlpT the mTMHs only insert efficiently when the entire subsequent helix is present, see Table 2. When either the positively charged residues or the hydrophobic part of the subsequent helix is absent the mTMHs are not recognized. This means that the positive charged residues located at the N-termini of a subsequent helix cause an orientational preference that then allows the mTMHs to be recognized by the translocon.

**Table 2:** Translocon recognition efficiency (Ins.Eff.) of marginally hydrophobic helices with and without additional arginines and subsequent helices. For EmrD the inclusion of flanks increases the number of positive charges on the outside loop, which causes it to insert worse. Inclusion of two arginines from *TMH*_*sub*_ does not change insertion efficiency much. EmrD mTMH2 is only inserted (40%) when T*MH*_*sub*_ is included. For GlpT the inclusion of flanks provides the outside loop with three positive charges and the inside loop with two. Addition of the arginine(s) from *TMH*_*sub*_ does not change this and in order to insert up to 37%, *TMH*_*sub*_ has to be included. For AcrB inclusion of the arginine residues of *TMH*_*sub*_ gives the inside loop six arginines in total against one on the outside loop, which brings the membrane integration up to 80% compared to 86% when *TMH*_*sub*_ is present.

| Name | Position | Included fragments | Ins.Eff. (%) | $\Delta KR$ |
| --- | --- | --- | --- | --- |
| EmrD H2F | 36-73 | Helix 2 + Flank | 1 | -1 |
| EmrD H2FR | 36-73 | Helix 2 + Flank + 2xArginines | 8 | 1 |
| EmrD H2-3 | 36-XX | Helix 2 + Flank + Helix 3 | 40 | 1 |
| GlpT H8F | 278-321 | Helix 7 + Flank | 8 | -1 |
| GlpT H8FR | 278-321 | Helix 7 + Flank + 1xArginine | 8 | 0 |
| GlpT H8-9 | 278-321 | Helix 7 + Flank + Helix 8 | 37 | 0 |
| AcrB H9-10F | 894-973 | Helix 9 + Helix 10 + Flank | 35 | 2 |
| AcrB H9-10FR | 894-973 | Helix 9 + Helix 10 + Flank + 3xArginines | 80 | 5 |
| AcrB H9-11 | 894-973 | Helix 9 + Helix 10 + Helix 11 | 86 | 5 |

In contrast, for AcrB mTMH10 inclusion of the six most N-terminal residues of the subsequent helix is sufficient to ensure membrane integration. These six residues include three arginines, and increases the total number of positively charged residues in the inside loop from three to six, see Table 2. Meanwhile, when the entire subsequent helix is present, these arginines are replaceable; not even a triple isoleucine mutant reduces the insertion of the mTMH. This further strengthens the observation that the positive-inside rule, caused by the positively charged residues in the loop, becomes more efficient when it is used to cause an oriental preference of a subsequent transmembrane helix.

### 4.1 The last and first helices in transmembrane proteins are more hydrophobic

Next, we examined if the presence of marginally hydrophobic helices correlates with positive charges in the loops. It could be expected that marginally hydrophobic helices would have a stronger positive bias in its surrounding loops than other helices. The positive inside bias is not significantly stronger for marginally hydrophobic helices in comparison to other helices, see Figure 6, nor is it significantly different in the preceding or subsequent helices of the mTMHs. This only demonstrates that the orientations of marginally hydrophobic, as well as other, helices are also affected by the positive inside rule.

**Figure 6:**
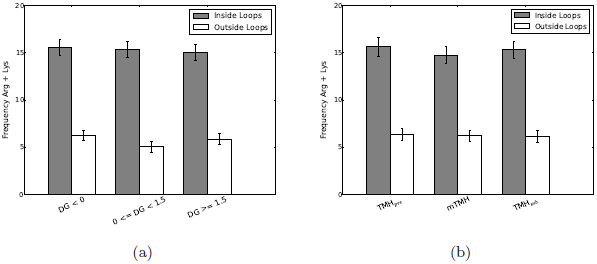
Frequency of positively charged amino acids in loops flanking transmembrane helices. a) Transmembrane helices sorted by hydrophobicity. b) Grouped into *TMH*_*pre*_, mTMH, and *TMH*_*sub*_. There are no significant differences for any of the groups.

To what extent is the orientation of subsequent helix crucial for the translocon recognition of marginally hydrophobic helices? A possibility to analyze this is to separate the last (and first) helix in multi-spanning transmembrane proteins, as these helices do not have any subsequent (or preceding) transmembrane helices. The positive inside bias is not significantly enhanced for the last (and first) helix in multi-spanning membrane proteins, see Figure 7A. However, the last helix is on average more hydrophobic than central transmembrane helices, see Figure 7B. The average hydrophobicity of the last helix is actually very similar to what is observed for the first helix in a transmembrane protein. It has been argued that the first helix needs to be more hydrophobic for proper membrane targeting^3^. Now it also appears as if the last helix needs to be sufficient hydrophobic for the positive inside rule to properly orient transmembrane helices.

**Figure 7:**
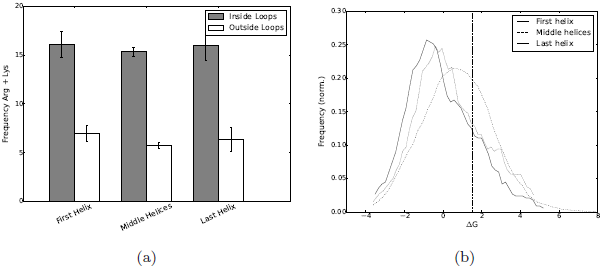
Distribution of positively charged residues in loops surrounding transmembrane helices. Here, only proteins containing three or more TMHs were considered. a) Frequency of positive charged residues in loops surrounding the first, last and internal helices. b) Hydrophobicity of transmembrane helices depending on the position in the protein. The curve is smoothed using a running average.

## 5 Conclusions

Here, we show that marginally hydrophobic helices in EmrD, GlpT and AcrB do not depend on specific interactions with their subsequent transmembrane helices. Instead, positively charged residues are important for the integration of the marginally hydrophobic helices. However, at least in EmrD and GlpT this effect is not directly mediated by the positive inside rule, but is rather an indirect effect, by creating an orientational preference of the subsequent transmembrane helices. The orientational effect of positively charged amino acids is thus enhanced when the residues are followed by a transmembrane helix. Supporting the importance of this orientational effect, we observe that marginally hydrophobic helices are rare as first or terminal helices of multi-spanning transmembrane proteins.

## Acknowledgments

This work was supported by grants from the Swedish Research Council (VR-NT 2009-5072, 2012-5046, VR-M 2010-3555), SSF, the Foundation for Strategic Research, Science for Life Laboratory, Swedish E-science Research Center, the EU 7'th Framework Program is gratefully acknowledged for support to the EDICT project, contract No: FP7-HEALTH-F4-2007-201924. The microsomes were gratefully given to us by Bernhard Dobberstein.

## 6 Material and methods

### 6.1 Enzymes and chemicals

Unless stated otherwise, chemicals were from Sigma-Aldrich (St. Louis, MO). Oligonucleotides were obtained from MWG Biotech AG (Ebersberg, Germany). Restriction enzymes were from Fermentas (Burlington, Ontario, Canada) and Phusion DNA polymerase from Finnzymes OY (Espoo, Finland). QuikChangeTM Site-Directed Mutagenesis Kit from Stratagene (La Jolla, CA). The plasmid pGEM-1 and the TNT SP6 Quick Coupled Transcription/Translation System were from Promega Biotech AB (Madison, WI). [^35^S]-methionine was from PerkinElmer (Boston, MA). Washed dog pancreatic microsomes were obtained from tRNAprobes (College Station, Texas). The E.Z.N.A Plasmid Purification, Cycle Pure and Gel Extraction kits from Omega Bio-Tek (Norcross, GA) were used during post-PCR manipulation and amplification of DNA

### 6.2 Introducing mTMH with neighboring helices into Leader peptidase host protein

The *lepB* gene had previously been introduced into the pGEM-1 vector, under the control of an SP6-promoter and with the context 5’ of the initiator codon changed to a Kozak consensus sequence^14^. To allow Lep to ”host” other protein segments, SpeI and KpnI restriction recognition sites had been introduced in the sequence encoding either the middle of the P2-domain (Lep-I) or on either side of TMH2 (Lep-II). The N-terminal tail of Lep-II had been lengthened to allow efficient glycosylation of the upstream glycosylation site^15^. Double-stranded oligonucleotides complementary to the 5’ and 3’ ends of the selected parts of AcrB, EmrD and GlpT were designed and used to amplify the mTMH together with its neighboring helices. The PCR products were cleaved with restriction enzymes SpeI and KpnI, separated on agarose gel and purified. These segments where then introduced into the lepB gene.

### 6.3 Engineering glycosylation sites as topology reporters

Native glycosylation sites NX[S/T]^16^ had been eliminated by substitution to QX[S/T] (where X can be any amino acid except for proline^17^) and new glycosylation acceptor sites had been engineered upstream of the SpeI-site and downstream of the KpnI-site, as well as within the introduced segments. For EmrD and GlpT constructs the first glycosylation site (NST) occurs after the second Lep-helix, the second between the mTMH and the subsequent helix (NRT), the third just after the insert (NAT) and final close to the C-terminal end (NST), see Figure 1). For AcrB construct, the first glycosylation site is at the N-terminus of the construct (NST), the second between the mTMH and the subsequent helix (NMT) and the third directly after the insert (NST) and one towards the C-terminal (NKT), see Figure 1. The glycosylation sites between mTMH and the subsequent helix are within a short loop and will only be fully glycosylated if none of the tested helices are inserted into the membrane.

### 6.4 Isoleucine and alanine scans

Point mutations to isoleucine (for AcrB and EmrD) or alanine (for GlpT) were performed by PCR using either QuickChangeTM Site-Directed Mutagenesis Kit or Phusion DNA polymerase. In order to avoid cleavage by signal peptidase, isoleucine was chosen as the substituting amino acid for EmrD and AcrB. In addition, the C-terminal of the EmrD-segment was modified, V89AVTT→I89IVYY, annotation according to native EmrD for the same reason. All DNA modifications were confirmed by sequencing of plasmid DNA at Eurofins MWG Operon. A few constructs failed to express and were consequently not included in this work.

### 6.5 Expression in vitro

Constructs in pGEM-1 were transcribed and translated in TNT SP6 Quick Coupled System from Promega. 100 ng DNA, 0.5 [^35^S]-methionine (5*μ*Ci) and dog pancreas rough microsomes were added to 5*μ*l of lysate at the start of the reaction and the samples were incubated at 30 °C for 90 min.

Two different batches of microsomes were used. The older batch was obtained from Bernhard Dobberstein, and the newer batch consisted of microsomes that had been column washed and purchased from tRNA Probes. In this study some systematic differences were observed between these two batches. Therefore, any mutant constructs should always be compared to the respetive wild type construct expressed with the same microsome batch. Hence all figures show the results for substitution scans together with those of the wild type construct. When a sub figure contains data obtained by two different microsome batches a star (^*^) will be used to denote data obtained with the new batch of microsomes. In both cases, the amount of microsomes was adjusted to guarantee 80% targeting. The microsome membranes harbor the protein complex Oligosaccharyl transferase (OST), with an enzyme with its active site on the lumen of the microsomes that will attach a glycan on the QX[S/T] site. Since glycosylation can only occur on the lumenal side, glycosylation can be used as a topology marker. Each glycan will add 2 kDa to the molecular weight and hence the differently glycosylated proteins can be separated on SDS-PAGE, see Figure 1.

### 6.6 Interpretation of differently glycosylated species

In EmrD and GlpT constructs the first glycosylation site occurs after the second Lep-helix. Both Lep helices are hydrophobic and will integrate into the membrane with N-terminus in the lumen. Because of this, the mTMH will end up in the lumen or insert into the membrane with N-terminus towards lumen. This is also true for the first Lep helix in LepII system used for AcrB constructs.

Previous studies ^3^ have shown that the AcrB, EmrD and GlpT *TMH*_*sub*_ and *TMH*_*pre*_ all are hydrophobic and are recognized to 82-100% by the translocon. In contast, the three mTMHs are only recognized to 8-13%. Hence, the probability to find a construct with the mTMH and not the *TMH*_*sub*_ inserted is low.

As described in Figure 1 different topologies should be possible to distnguish dependent on the number of glycosylation sites. For all constructs the tripple glycosylated form represent the insertion of the entire construct, while double glycosylated sites should in theory represent the constructs where mTMH is not inserted. However, the double glycosylated form is rarely observed, but instead the single glycosylated form is common for most constructs.

In this study, as well in our earlier study^3^ we have interpreted the singly glycosylated band as representing the toplogy where only the mTMH is not inserted. The short loop between mTMH and *TMH*_*sub*_ cause inefficient glyco-sylation of this glycosylation site, resulting in that this topology might be either singly or doubly glycosylated. The loop is only a few amino acids long and it is known that short loops are often inefficiently glycosylated^15,18^.

We have previously shown that none of the mTMHs studied here, are inserted into the membrane to any larger degree (¡13%). Meanwhile, all *TMH*_*sub*_ are hydrophobic and readily recognized by the translocon (80-100%)^3^. Further, Δ*G*_*pred*_ for any variant of the *TMH*_*sub*_ is much lower, -2.65 - +0.34, than for the mTMHs (+2.40 - 3.46), indicating that the topology III is very unlikely to occur. For EmrD and GlpT topology IV has not been observed, whereas a few AcrB constructs gave rise to a low amounts of topology V.

### 6.7 Data analysis

The proteins were visualized with a Fuji FLA-3000 phosphoimager (Fujifilm,

Tokyo, Japan) with the Image Reader V1.8J/Image Gauge V 3.45 software (Fujifilm). The MultiGauge software was used to create one-dimensional intensity profile for each lane on the gels. These profiles were then analyzed using the multi-Gaussian fit program from the Qtiplot software package (http://www.qtiplot.ro/), and the peak areas of the glycosylated protein bands in the profile were calculated. Each peak represents the protein with 0, 1, 2, 3 or 4 glycan molecules.

Hence, the frequency of the different topologies for each construct can be deduced.

### 6.8 Statistical analysis of mTMHs

For the statistical analysis the 289 proteins with experimentally verified topologies from an earlier study were used^19^. All these proteins have a sequence identity of less than 20% to any other member of the dataset. It is difficult to exactly identify the membrane boarders and length of a transmembrane helix^11^ and it has been shown that transmembrane helices might shift in the membrane region after recognition by the translocon^12^. Therefore, the membrane-spanning region of each transmembrane helix is defined as the most hydrophobic twenty-one residue long segment shifted by maximum 5 residues from the experimentally detected transmembrane region. The hydrophobicity of each helix was then estimated using the ΔG predictor^1^.

The standard errors, represented as error bars in Figures 6 and 7 were calculated using equation (1), where *σ* is the standard deviation and *n* is the number of samples.

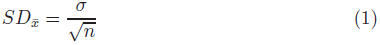

## References

[1] Hessa, T., Meindl-Beinker, N., Bernsel, A., Kim, H., Sato, Y., Lerch-Bader, M., Nilsson, I., White, S., and von Heijne, G. Molecular code for transmembrane-helix recognition by the Sec61 translocon. Nature 450(450):1026–1030, Dec, 2007.

[2] Lundin, C., Kim, H., Nilsson, I., White, S., and von Heijne, G. Molecular code for protein insertion in the endoplasmic reticulum membrane is similar for N(in)-C(out) and N(out)-C(in) transmembrane helices. Proc Natl Acad Sci USA 105(41):15702–15707, Oct, 2008.

[3] Hedin, L., Ojemalm, K., Bernsel, A., Hennerdal, A., Illergard, K., Enquist, K., Kauko, A., Cristobal, S., von Heijne, G., Lerch-Bader, M., Nilsson, I., Elofsson, A. Membrane insertion of marginally hydrophobic transmembrane helices depends on sequence context. J Mol Biol 396(1):221–229, Feb, 2010.

[4] Sadlish, H. Skach, W. Biogenesis of CFTR and other polytopic membrane proteins: new roles for the ribosome-translocon complex. J Membr Biol 202(3):115–126, Dec, 2004.

[5] Pitonzo, D. Skach, W. Molecular mechanisms of aquaporin biogenesis by the endoplasmic reticulum Sec61 translocon. Biochim Biophys Acta 1758(8):976–988, Aug, 2006.

[6] Sato, Y., Sakaguchi, M., Goshima, S., Nakamura, T., Uozumi, N. Integration of Shaker-type K+ channel, KAT1, into the endoplasmic reticulum membrane: synergistic insertion of voltage-sensing segments, S3-S4, and independent insertion of pore-forming segments, S5-P-S6. Proc Natl Acad Sci U S A 99(1):60–65, Jan, 2002.

[7] Meindl-Beinker, N., Lundin, C., Nilsson, I., White, S., and von Heijne, G. Asn- and Asp-mediated interactions between transmembrane helices during translocon-mediated membrane protein assembly. EMBO Rep 7(11):1111–1116, Nov, 2006.

[8] Zhang, L., Sato, Y., Hessa, T., von Heijne, G., Lee, J., Kodama, I., Sakaguchi, M., Uozumi, N. Contribution of hydrophobic and electrostatic interactions to the membrane integration of the Shaker K+ channel voltage sensor domain. Proc Natl Acad Sci USA 104(20):8263–8268, May, 2007.

[9] Ojemalm, K., Halling, K., Nilsson, I., and von Heijne, G. Orientational preferences of neighboring helices can drive ER insertion of a marginally hydrophobic transmembrane helix. Mol Cell 45(4):529–540, Feb, 2012.

[10] Bano-Polo, M., Martinez-Gil, L., Wallner, B., Nieva, J., Elofsson, A., Mingarro, I. Charge Pair Interactions in Transmembrane Helices and Turn Propensity of the Connecting Sequence Promote Helical Hairpin Insertion. J Mol Biol 0, Dec, 2012.

[11] Papaloukas, C., Granseth, E., Viklund, H., and Elofsson, A. Estimating the length of transmembrane helices using Z-coordinate predictions. Protein Sci 17(2):271–278, Feb, 2008.

[12] Kauko, A., Hedin, L., Thebaud, E., Cristobal, S., Elofsson, A., and von Heijne, G. Repositioning of transmembrane alpha-helices during membrane protein folding. J Mol Biol 397(1):190–201, Mar, 2010.

[13] von Heijne, G. Gavel, Y. Topogenic signals in integral membrane proteins. Eur J Biochem 174(4):671–678, Jul, 1988.

[14] Kozak, M. Context effects and inefficient initiation at non-AUG codons in eucaryotic cell-free translation systems. Mol Cell Biol 9(11):5073–5080, Nov, 1989.

[15] Nilsson, I. and von Heijne, G. Determination of the distance between the oligosaccharyltransferase active site and the endoplasmic reticulum membrane. J Biol Chem 268(8):5798–5801, Mar, 1993.

[16] Silberstein, S. Gilmore, R. Biochemistry, molecular biology, and genetics of the oligosaccharyltransferase. FASEB J 10(8):849–858, Jun, 1996.

[17] Shakin-Eshleman, S., Spitalnik, S., Kasturi, L. The amino acid at the X position of an Asn-X-Ser sequon is an important determinant of N-linked core-glycosylation efficiency. J Biol Chem 271(11):6363–6366, Mar, 1996.

[18] Armulik, A., Nilsson, I., von Heijne, G., Johansson, S. Determination of the border between the transmembrane and cytoplasmic domains of human integrin subunits. J Biol Chem 274(52):37030–37034, Dec, 1999.

[19] Bernsel, A., Viklund, H., Falk, J., Lindahl, E., von Heijne, G., Elofsson, A. Prediction of membrane-protein topology from first principles. Proceedings of the National Academy of Sciences of the United States of America 105(20):7177–7181, May, 2008.

